# A multimodal representation learning platform for accurate molecular ADMET prediction

**DOI:** 10.64898/2026.08.24.746660

**Authors:** Zhensheng Luo, Dachen Huang, Yanruisheng Shao, Qinze Yu, Yu Li

## Abstract

**Motivation:** Accurate ADMET prediction is essential for prioritizing compounds before costly experimental validation, yet ADMET tasks are highly heterogeneous. Properties such as solubility, permeability, protein binding, clearance, transporter activity and toxicity are governed by different molecular signals, ranging from local functional groups and physicochemical descriptors to bonded topology and three-dimensional geometry. Consequently, a single molecular representation or backbone is unlikely to be optimal across all ADMET tasks.

**Results:** We present Trimole-Hybrid, a task-wise multimodal framework that addresses ADMET heterogeneity by selecting or combining predictors built from complementary molecular representations. Trimole-Hybrid constructs a candidate pool of SMILES-, graph-, geometry-sensitive EPT/3D- and chemical descriptor-based predictors. For each task, Trimole-Hybrid selects the best-performing predictor to obtain the final prediction. On 22 Therapeutics Data Commons ADMET benchmarks, Trimole-Hybrid exceeded the public TDC top-1 methods on 10 tasks and ranked within the top 10 for 21 tasks. Ablation studies confirmed the contribution of both complementary multimodal molecular representations and task-specific ensemble strategies. In two small-molecule case studies, Trimole-Hybrid shows sensitivity to changes in essential functional motifs, suggesting its ability to capture ADMET-relevant molecular substructures.

**Availability and implementation:** Source code, supplementary tables, and audit files are available at the project repository: https://github.com/dchen0212/trimole_hybrid.

**Supplementary information:** Supplementary data are available with the submitted manuscript.

## Introduction

Computational ADMET prediction helps screen drug candidates before costly experiments. A compound can bind its intended target and still fail because it is poorly absorbed, cleared too quickly, affected by transporters or associated with toxicity. ADMET models are useful when they help chemists rank compounds earlier and focus experiments on molecules with a better chance of progressing [24, 3, 26].

The difficulty is that ADMET is a collection of related but distinct assays, not a single prediction problem. The TDC ADMET benchmark includes small and large datasets, classification and regression tasks, and metrics such as AUROC, AUPRC, MAE and Spearman correlation [11]. The relevant chemistry also changes across assays: solubility, permeability, protein binding, metabolic stability and toxicity draw on different mixtures of polarity, hydrophobicity, size, charge, topology and three-dimensional structure. A representation that works well for one property may miss the information needed for another.

Current molecular learning methods capture different parts of this chemistry. Fingerprints and chemical descriptors remain useful, especially when datasets are small [15, 18, 26]. Deep models can learn from SMILES strings, molecular graphs, 3D conformations and pretrained molecular embeddings [30, 27, 20, 9, 31]. Shared benchmarks such as MoleculeNet and TDC make it possible to compare these model classes under common splits and metrics [29, 11, 7]. A recurring lesson from these comparisons is that no representation is uniformly best; performance depends on the task, the data size and the molecular information available to the model [30, 7].

Together, these differences create a benchmark problem as well as a modeling problem. ADMET results are often summarized as if one molecular backbone should dominate the full benchmark. For heterogeneous ADMET tasks, however, a single architecture name is less informative than the molecular view and model combination used for each task.

These differences motivate a multimodal candidate pool rather than a single input representation. Descriptor models retain explicit physicochemical and fragment-level priors. Graph and SMILES encoders capture bonded topology and recurring molecular motifs. EPT/3D-related models add geometry-sensitive information when conformation or steric effects matter.

Trimole-Hybrid targets this heterogeneity directly. It creates a pool of sequence, graph, geometry-sensitive and descriptor-based predictors, and assigns a final model setup separately for each ADMET task. Rather than forcing all tasks through one fused representation, it records which molecular information source, model family and prediction average were used for each property. In this sense, Trimole-Hybrid turns multimodal ADMET prediction from a search for one best backbone into a task-wise model-composition problem.

We tested this view on the 22-task TDC ADMET benchmark. The fixed task setups matched or exceeded the public TDC top-1 reference on 10 tasks and ranked in the top 10 on 21 tasks. Ablation experiments then showed that chemical descriptors, EPT/3D information, prediction averaging, seed averaging and task-wise model choice contributed to these scores. Finally, two molecule-level examples connected the benchmark to individual chemical structures: an AMES-positive benchmark molecule and an external Deutivacaftor/P-gp transporter case. In both examples, Trimole-Hybrid classified the original molecule correctly, and local sensitivity tests examined chemically plausible regions without treating them as causal pharmacophores. Together, these results support a practical reporting principle: ADMET benchmarks should report which molecular view and model combination worked for each task, not just a single model name.

## Materials and methods

### Datasets and evaluation

We tested Trimole-Hybrid on the 22 tasks in the TDC ADMET benchmark group. Each task corresponds to one dataset and one measured ADMET property. Together, the tasks cover five practical questions in early drug discovery: absorption, distribution, metabolism, excretion and toxicity. Absorption tasks measure solubility, barrier crossing and transporter-related processes. Distribution tasks describe where a compound is likely to go in the body. Metabolism tasks capture cytochrome P450 substrate or inhibition behavior. Excretion tasks measure clearance-related properties. Toxicity tasks cover mutagenicity, liver injury, hERG liability and acute toxicity.

All experiments used the official TDC/admet-group scaffold splits [11, 1]. The supplied training and validation partitions were used during model development, and the supplied test partition was used only for final scoring. The official task metrics were AUROC, AUPRC, mean absolute error (MAE) or Spearman correlation.

To compare tasks with different metric directions, we used a direction-normalized margin

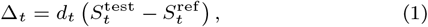

where 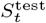 is the final Trimole-Hybrid test score, 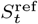 is the public TDC top-1 reference score, and *d*_*t*_ is the metric direction. For AU-ROC, AUPRC and Spearman correlation, *d*_*t*_ = 1; for MAE, *d*_*t*_ = −1. A positive Δ_*t*_ means that Trimole-Hybrid matched or exceeded the reference after accounting for metric direction. Unless stated otherwise, final benchmark scores are reported as five-run mean*±*standard deviation.

### Candidate models and model choice

Trimole-Hybrid uses a candidate pool rather than one fixed backbone. The pool contains four main molecular views. SMILES encoders provide sequence-level information from canonical molecular strings and capture recurring local motifs [6]. Graph encoders provide atom-bond connectivity through KPGT-style representations, preserving topology and substructure context [13]. EPT/3D-related models add geometry-sensitive information when shape, conformation or steric effects are relevant [9, 31, 21]. Chemical descriptor models use Morgan/ECFP-like fingerprints, RDKit descriptors and tabular chemical features to represent fragments and physicochemical properties [15, 19, 18].

The chemical feature branch trains lightweight models such as XG-Boost, ExtraTrees, random forests or logistic regression [5, 2, 17]. Prediction files can also be averaged across input types, scaffold-CV folds or random seeds [28, 8]. For each task, candidate ranking used the official validation set or scaffold-cross-validation folds made from the training/validation data. After a candidate was selected, its model type, averaging rule and seed set were fixed before official-test scoring. The separation prevents repeated test-set checking from influencing model choice [7, 25, 4]. Additional notation for input types, averaging and candidate ranking is provided in Supplementary Note 3, and the final predictor, file identifiers and selection record for each task are listed in Supplementary Tables S2, S13 and S15.

### Benchmark reporting and reference comparison

For each task, we recorded the model family, validation metric, chosen predictor, seed set, averaging rule, EPT/3D status and prediction-file identifiers. TDC top-1 references and top-*k* ranks were taken from the public leaderboard record available on 15 May 2026. The top-1 reference is the best public score in that leaderboard record for the same task and metric. Official test labels and leaderboard outcomes were not used to choose model families, averaging weights or seed sets. The metric-level comparison in Figure 2c used one public model per metric family when matched public scores were available under the same TDC split and metric. The comparisons were descriptive and were not used for model choice or rank assignment.

**Figure 1.**
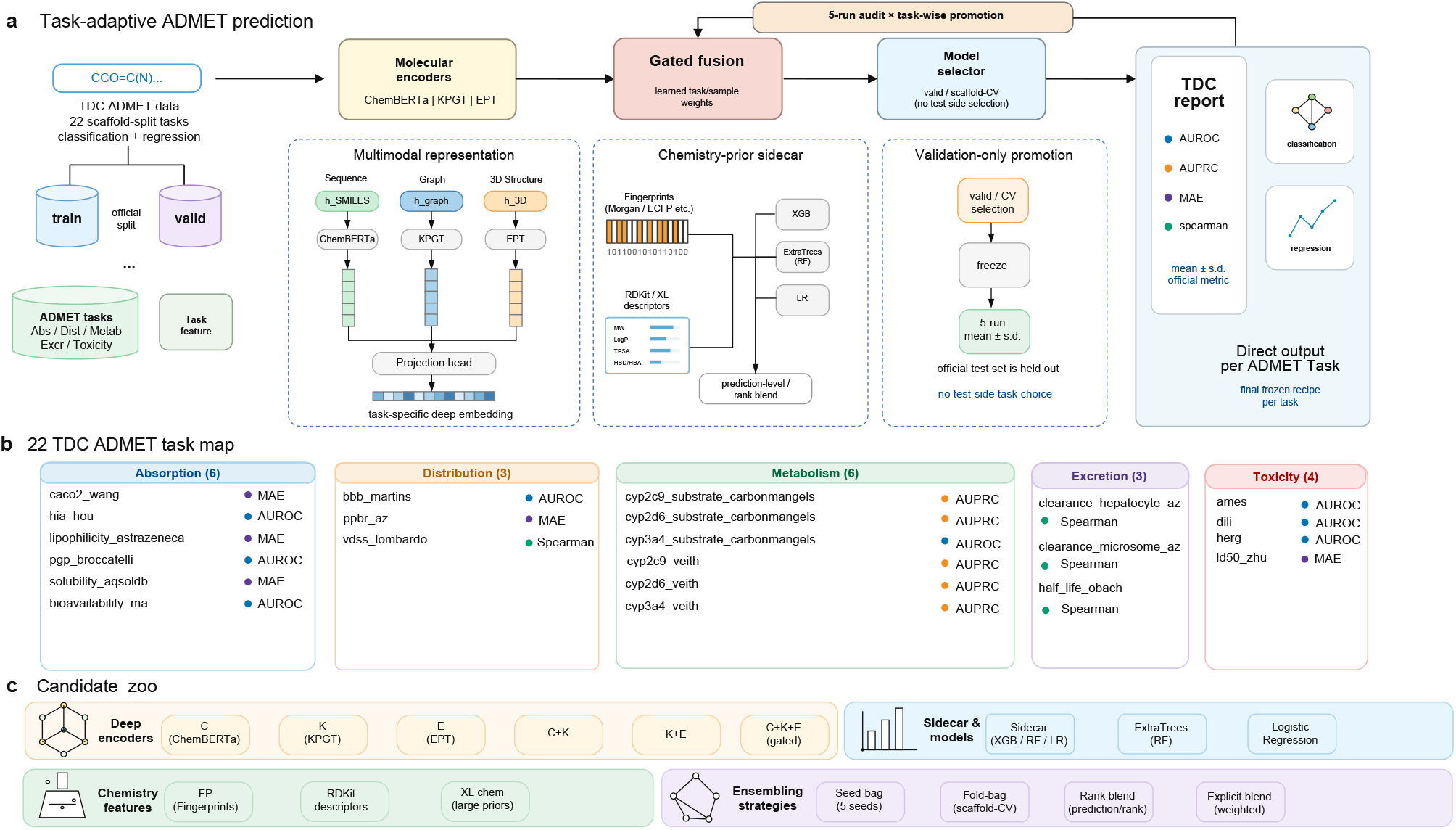
Overview of Trimole-Hybrid. **a**, Starting from the official TDC scaffold splits, each molecule is encoded with SMILES, graph, 3D-related and chemical descriptor features. Candidate predictors and prediction averages are chosen separately for each task, then the fixed task setup is evaluated on the official test split. **b**, The 22 TDC ADMET tasks grouped by ADMET category and official metric. **c**, Candidate model pool, including deep encoders, chemical descriptor models and prediction averaging methods used to build the final task-specific predictors.

**Figure 2.**
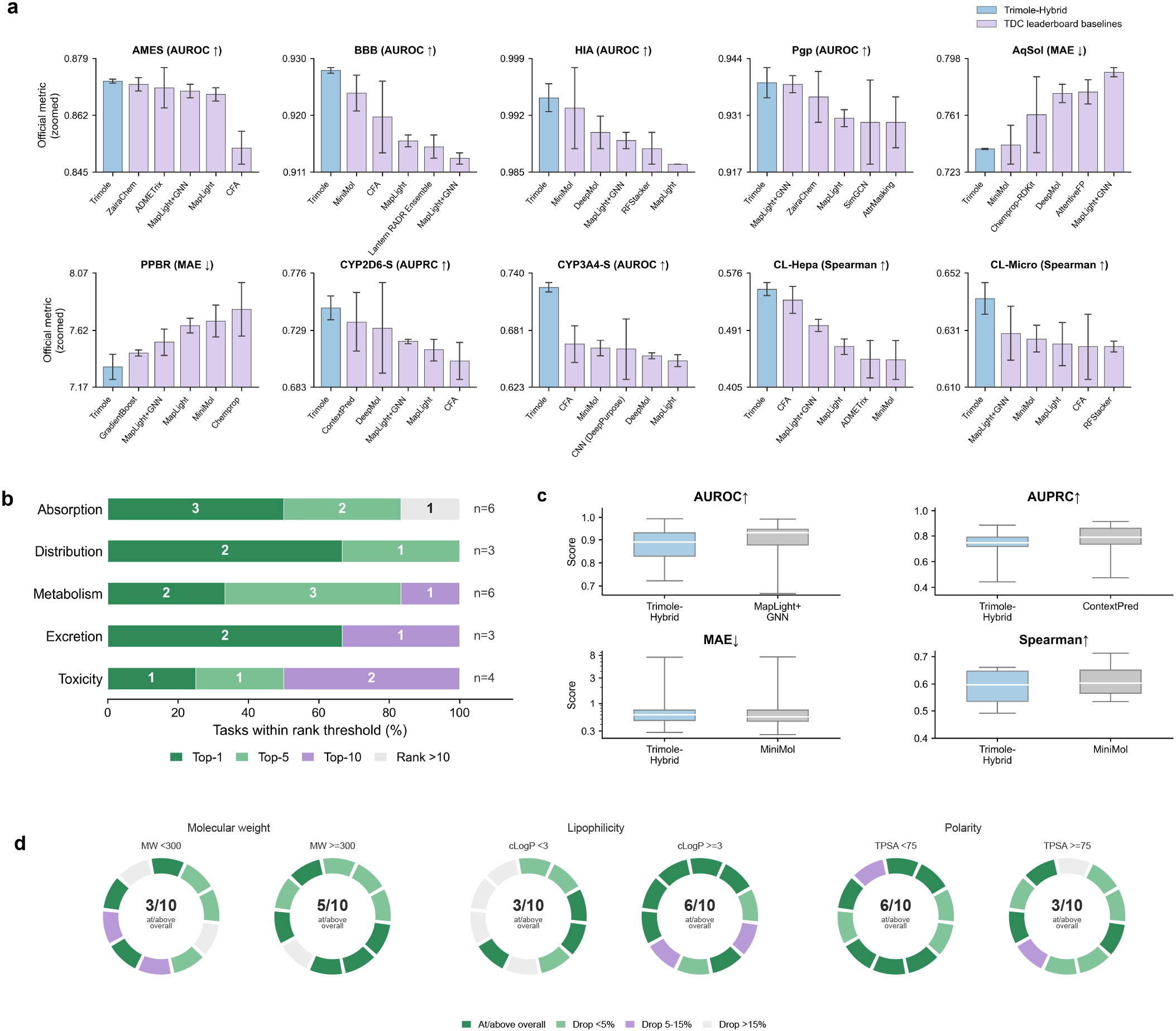
Task-level and grouped benchmark results for Trimole-Hybrid on the 22-task TDC ADMET benchmark. **a**, Official scores for the 10 tasks where Trimole-Hybrid matched or exceeded the public TDC top-1 reference, shown against public leaderboard baselines under the same task metric. **b**, ADMET-wise rank coverage. Each horizontal bar summarizes all tasks within one ADMET category and shows the fraction of tasks that reached the public top-1 reference, remained within the top-5 range, remained within the top-10 range, or fell outside the top 10. **c**, Within-metric normalized score distributions for Trimole-Hybrid and a matched public comparison model. **d**, Molecule-level subgroup analysis for the 10 top-reference tasks. Official test molecules were split by molecular weight, cLogP and TPSA, and the official metric was recomputed within each subgroup. Each ring contains the 10 tasks. The centre value gives the number of tasks whose subgroup score stayed at or above the overall task score; the colour of each segment shows whether the corresponding subgroup showed no drop, a small drop, a moderate drop or a large drop.

## Results

Trimole-Hybrid was evaluated in three steps. First, we tested whether task-wise model composition could compete with public TDC ADMET references. Second, we checked whether the gains were concentrated in particular ADMET categories, metric types or molecule-property ranges. Third, we used ablations and molecule-level sensitivity tests to examine which parts of the system mattered.

### Benchmark performance across ADMET tasks

Table 1 reports the main benchmark result: the official-test score for all 22 tasks, the public top-1 reference, the direction-normalized margin and the rank range. Figure 2 summarizes the same evidence from four angles: task-level scores, ADMET-category rank coverage, metric-level score distributions and molecular-property subgroup checks.

**Table 1.** TDC ADMET 22-task benchmark results of Trimole-Hybrid. Scores are five-run mean *±*s.d. values from the final fixed model setup. Positive margins indicate matching or exceeding the public TDC top-1 reference after accounting for metric direction.

| Category | Dataset | Size | Metric | Dir. | Trimole-Hybrid | Top-1 ref. | Margin | Rank |
| --- | --- | --- | --- | --- | --- | --- | --- | --- |
| Absorption | TDC.Bioav | 640 | AUROC | ↑ | 0.722248±0.004048 | 0.942 | -0.219752 | Top-21 |
| Absorption | TDC.Caco2 | 906 | MAE | ↓ | 0.283647±0.002290 | 0.256 | -0.027647 | Top-5 |
| Absorption | TDC.HIA | 578 | AUROC | ↑ | 0.994280±0.001736 | 0.993 | 0.001280 | Top-1 |
| Absorption | TDC.Lipo | 4,200 | MAE | ↓ | 0.472895±0.006499 | 0.456 | -0.016895 | Top-4 |
| Absorption | TDC.Pgp | 1,212 | AUROC | ↑ | 0.938363±0.003565 | 0.938 | 0.000363 | Top-1 |
| Absorption | TDC.AqSol | 9,982 | MAE | ↓ | 0.738443±0.000456 | 0.741 | 0.002557 | Top-1 |
| Distribution | TDC.BBB | 1,975 | AUROC | ↑ | 0.927779±0.000477 | 0.924 | 0.003779 | Top-1 |
| Distribution | TDC.PPBR | 1,797 | MAE | ↓ | 7.330617±0.098555 | 7.440 | 0.109383 | Top-1 |
| Distribution | TDC.VD | 1,130 | Spearman | ↑ | 0.660995±0.004668 | 0.713 | -0.052005 | Top-3 |
| Metabolism | TDC.CYP2C9-S | 666 | AUPRC | ↑ | 0.443370±0.023358 | 0.474 | -0.030630 | Top-2 |
| Metabolism | TDC.CYP2C9-I | 12,092 | AUPRC | ↑ | 0.790586±0.002608 | 0.859 | -0.068414 | Top-8 |
| Metabolism | TDC.CYP2D6-S | 664 | AUPRC | ↑ | 0.747531±0.009610 | 0.736 | 0.011531 | Top-1 |
| Metabolism | TDC.CYP2D6-I | 13,130 | AUPRC | ↑ | 0.719172±0.005514 | 0.790 | -0.070828 | Top-4 |
| Metabolism | TDC.CYP3A4-S | 667 | AUROC | ↑ | 0.725249±0.004760 | 0.667 | 0.058249 | Top-1 |
| Metabolism | TDC.CYP3A4-I | 12,328 | AUPRC | ↑ | 0.884121±0.003206 | 0.916 | -0.031879 | Top-4 |
| Excretion | TDC.CL-Hepa | 1,020 | Spearman | ↑ | 0.552040±0.009650 | 0.536 | 0.016040 | Top-1 |
| Excretion | TDC.CL-Micro | 1,102 | Spearman | ↑ | 0.642814±0.005773 | 0.630 | 0.012814 | Top-1 |
| Excretion | TDC.Half_Life | 667 | Spearman | ↑ | 0.491390±0.009785 | 0.576 | -0.084610 | Top-7 |
| Toxicity | TDC.AMES | 7,255 | AUROC | ↑ | 0.871945±0.000583 | 0.871 | 0.000945 | Top-1 |
| Toxicity | TDC.DILI | 475 | AUROC | ↑ | 0.908609±0.005520 | 0.956 | -0.047391 | Top-6 |
| Toxicity | TDC.hERG | 648 | AUROC | ↑ | 0.863358±0.002334 | 0.880 | -0.016642 | Top-6 |
| Toxicity | TDC.LD50 | 7,385 | MAE | ↓ | 0.603071±0.003837 | 0.552 | -0.051071 | Top-5 |

Trimole-Hybrid matched or exceeded the public TDC top-1 reference on 10 tasks: AMES, BBB, clearance hepatocyte, clearance microsome, CYP2D6 substrate, CYP3A4 substrate, HIA, P-gp, PPBR and solubility. It ranked in the top 3 on 12 tasks, in the top 5 on 17 tasks and in the top 10 on 21 of 22 tasks. These gains were not confined to a single assay family.

The largest positive margins were observed for PPBR, CYP3A4 substrate, clearance hepatocyte, clearance microsome and CYP2D6 substrate. These improvements did not come from one repeated predictor. The top-reference tasks used different final setups, including chemical descriptor models, KPGT-based seed or prediction averaging, transfer-based XGBoost and rank-based averaging. The variation supports the central design choice: model choice should be made separately for each ADMET property. Negative-margin tasks, including bioavailability, lipophilicity, VDss, DILI, hERG and LD50, identify cases where the current candidate pool still lacks enough assay-specific signal.

### Performance stratified by ADMET category and metric type

A strong aggregate result could still be misleading if the gains were concentrated in one ADMET class. Figure 2b shows that this was not the case, but the coverage was uneven. Distribution and excretion had the clearest top-reference coverage. Absorption was mixed because its tasks measure different phenomena, from solubility and intestinal uptake to transporter behavior and bioavailability. Metabolism and toxicity had fewer top-reference matches, but most of these tasks still remained within the top-10 range. This pattern was not explained by dataset size alone: some small substrate datasets reached the top reference, whereas several much larger CYP inhibition or toxicity datasets did not. The harder tasks therefore appear to reflect endpoint-specific assay noise, class balance, metric sensitivity and public-baseline strength, not only the number of training molecules.

The apparent advantage also depended on the evaluation metric. Figure 2c separates the comparison by metric family. AUROC tasks mainly tested binary ranking over the whole test set and were competitive with MapLight + GNN [16]. AUPRC tasks were more sensitive to positive-class recovery, and the clearest gains appeared in substrate tasks. MAE tasks compared absolute prediction error on continuous properties; Trimole-Hybrid was strong for solubility and PPBR but did not dominate every regression endpoint. Spearman tasks measured rank ordering rather than absolute value, which explains why clearance and VDss behaved differently from MAE tasks. This break-down is descriptive and helps interpret where the method works best; Table 1 remains the formal leaderboard comparison.

We also tested whether the top-reference results were stable across simple chemical regimes. Official test molecules were split within each task by molecular weight, cLogP and TPSA, and the official metric was recomputed inside each subgroup. The analysis used the same fixed test predictions and did not retrain models or choose new task setups. Because matched per-molecule baseline predictions were unavailable, this is an internal robustness check rather than a direct comparison with public methods. The cLogP *≥* 3 and TPSA *<* 75 groups stayed at or above the overall task score in 6 of 10 tasks, whereas the MW *<* 300, cLogP *<* 3 and TPSA *≥* 75 groups reached that level in 3 of 10 tasks. Thus, the selected predictors retained signal across several molecule-property ranges, but the subgroup analysis also shows where additional chemistry-specific modeling may be needed.

### Ablation study

The ablation analysis tested whether the final scores came from one component or from several parts of the system. We compared the full setup with five restricted variants: no task-wise selection, no chemical descriptor models, no EPT/3D-containing models, no prediction-level averaging and no seed averaging. For each ablation, we removed one component, applied the same candidate-ranking procedure where applicable and then reported the official-test score. The single-seed comparison used the closest available single-seed model. Each ablation covers all 22 tasks in the checked summary. The comparison measures the value of each component under the documented workflow; it does not search for a new hand-picked alternative after inspecting test labels.

For an ablation variant *a*, the task-level effect was measured as

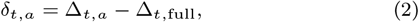

where Δ_*t*,full_ and Δ_*t,a*_ are the direction-normalized margins of the full and ablated versions. Negative values mean that the ablated version performed worse than the full method. Margin counts are score-versus-reference counts, not official leaderboard ranks, because the ablation versions were not submitted as leaderboard entries and do not have complete rank records.

Removing the chemical descriptor models caused the largest loss, while smaller but consistent drops followed removal of EPT/3D-containing models, prediction averaging, seed averaging or task-wise model choice. Fingerprints, RDKit descriptors and tabular models therefore provided signal that the deep molecular encoders did not fully replace. The ablation pattern indicates that the benchmark scores were not driven by a single backbone, but by combining assay-specific representation choice with conventional chemical priors.

As an additional diagnostic, a naive MLP late-fusion control trained on the same fixed candidate-model prediction files performed below the final setup on all 22 tasks (Figure 3e). Thus, simply adding a generic learned combiner did not replace task-wise model choice.

**Figure 3.**
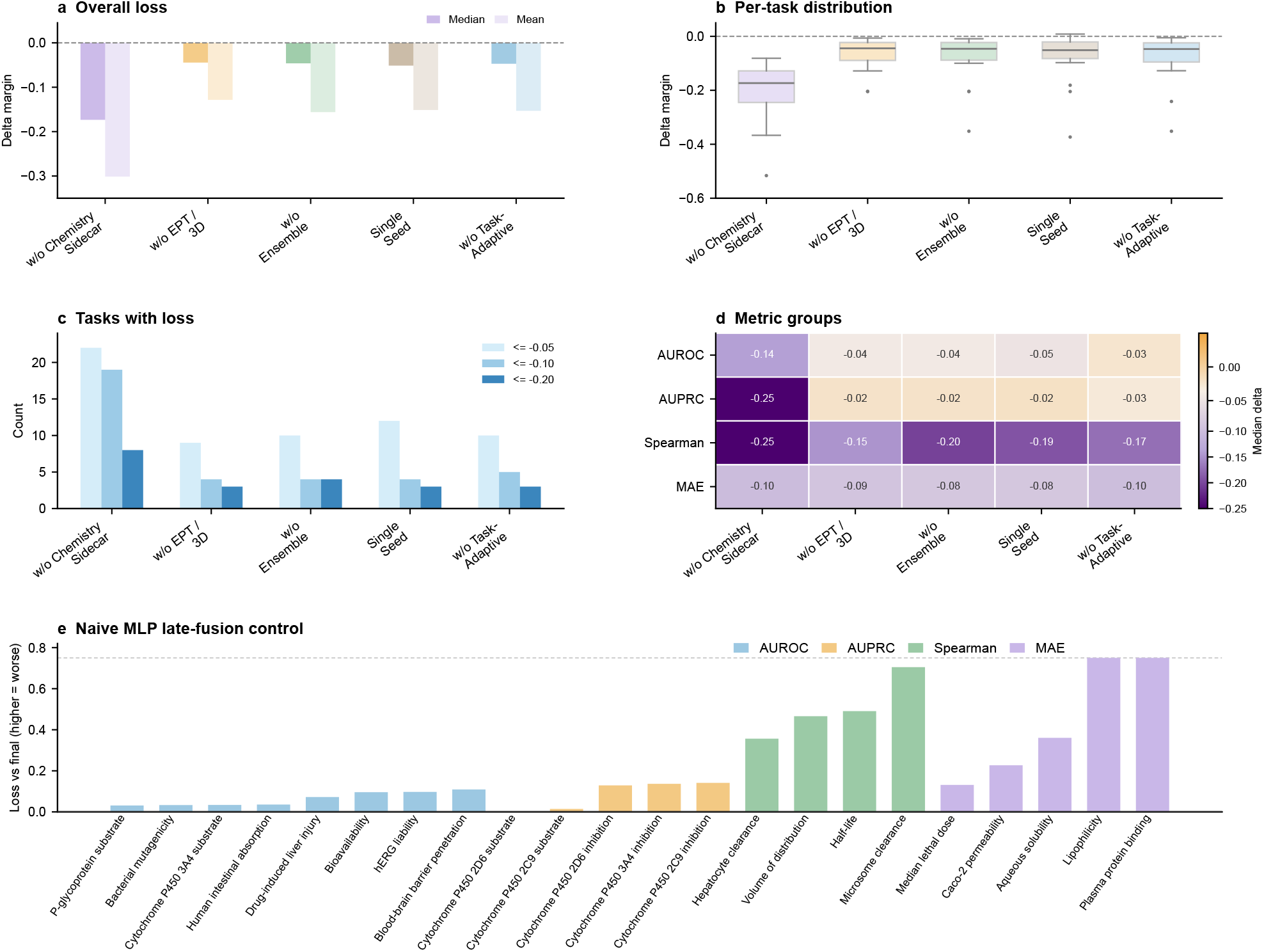
Formal ablation analysis shows which parts of Trimole-Hybrid affect performance. **a**, Mean and median changes in direction-normalized margin after removing each component. **b**, Per-task distribution of ablation-induced margin changes. **c**, Number of tasks exceeding predefined loss thresholds. **d**, Median margin changes by metric group. **e**, A naive MLP was trained to combine the same cached candidate-model predictions for each task. It performed worse than the final Trimole-Hybrid setup on all 22 tasks, showing that generic learned fusion did not replace task-wise model choice. Ablations and the MLP control are task-level comparisons for interpretation, not independent leaderboard submissions.

### Molecule-level sensitivity case studies

To complement the aggregate benchmarks, we used two individual molecules to check whether the task-specific predictors behaved sensibly at the compound level. The examples were an AMES-positive TDC test molecule and an external Deutivacaftor/P-gp substrate case from drug-label evidence. They were not used to choose models, promote tasks, design ablations or compare leader-board ranks. For each example, Figure 4 compares the MiniMol-style score, the Trimole-Hybrid score for the original molecule and the Trimole-Hybrid score after a matched local perturbation. The external molecule was checked against the relevant TDC splits and did not overlap with the corresponding training, validation or test split. Local molecule changes were evaluated either with the exact task pipeline or with a reconstructed task-specific proxy when exact full-model replay for arbitrary SMILES was not available. The highlighted regions mark tested perturbation sites, not experimentally validated pharmacophores.

**Figure 4.**
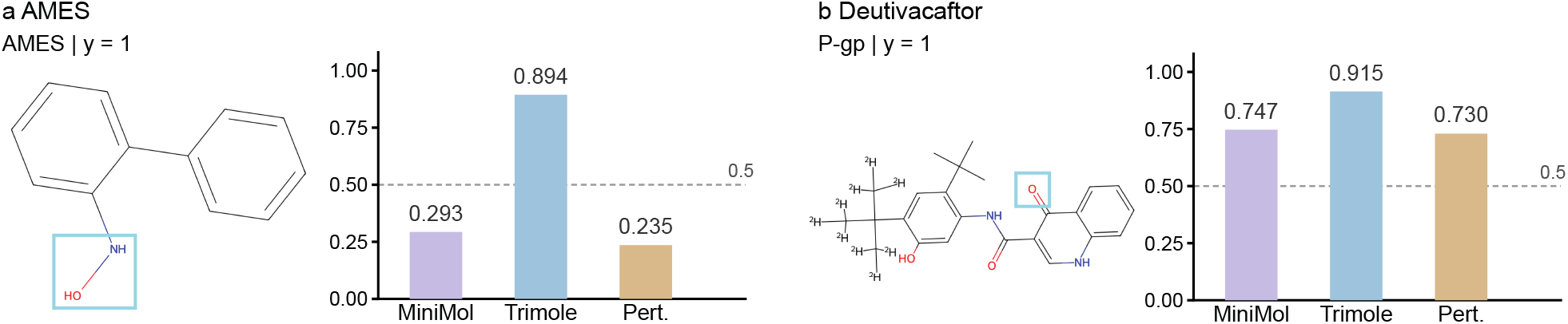
Molecule-level sensitivity case studies. **a**, AMES test_297 example. Bars show the MiniMol-style score, the materialized Trimole-Hybrid score for the original molecule and the perturbation score. The perturbation test uses a reconstructed model and is not exact full-model attribution. **b**, Deutivacaftor/P-gp example evaluated with the exact P-gp pipeline. Orange boxes mark the regions changed in the sensitivity test. They are not experimentally validated causal pharmacophores.

The two examples address complementary questions. The AMES case examines a benchmark molecule with a plausible mutagenic group, whereas the Deutivacaftor case tests a drug-relevant molecule outside the benchmark. Deutivacaftor is part of the FDA-approved cystic fibrosis drug Alyftrek, and transporter behavior can affect both drug exposure and drug–drug interaction risk [23].

AMES test_297 (*y* = 1; SMILES `ONc1ccccc1-c1ccccc1`) is not a marketed drug, but it provides a chemically interpretable mutagenicity example. The molecule contains a highlighted N-hydroxy/aniline-like region. N-hydroxylated aromatic amine motifs are known in genotoxicity chemistry because they can form reactive species linked to DNA damage and positive bacterial mutagenicity assays [12, 22]. The MiniMol-style prediction was low (0.293194), whereas the Trimole-Hybrid prediction was high (0.894085). Exact full-model replay for arbitrary AMES SMILES was not available, so we tested the local molecule change with a reconstructed fingerprint-XGBoost model for this task. Replacing the highlighted region reduced the reconstructed score from 0.894710 *±* 0.008471 to 0.235478 *±* 0.009304. The decrease is consistent with the high final AMES score, but it is not exact attribution for the full Trimole-Hybrid model.

Deutivacaftor provides an external pharmacology example. It is a CFTR potentiator used in Alyftrek. For such a drug, exposure and transporter handling matter in addition to target activity. The prescribing information discusses Deutivacaftor in the P-gp/BCRP transporter context [23]. In the external P-gp substrate case (*y* = 1), the MiniMol-style prediction was 0.746979 *±* 0.064799, whereas the exact Trimole-Hybrid replay gave 0.914545 *±* 0.021395. We tested the highlighted oxygen-containing linker because this region can affect polarity, hydrogen-bond acceptor count and conformation. Replacing this region with a methylene group reduced the exact P-gp score to 0.729853 *±* 0.149185 while keeping the prediction positive. The decrease indicates sensitivity to this polar linker, but does not prove that the linker is a causal pharmacophore [14].

## Discussion

The benchmark and ablation results support a task-wise view of ADMET prediction. Trimole-Hybrid matched or exceeded the public TDC top-1 reference on 10 tasks, but those tasks did not use one common model setup. This agrees with the chemistry of ADMET assays: clearance, permeability, transporter activity, protein binding and toxicity depend on different molecular features [30, 9, 31, 7]. It is therefore unlikely that a single molecular representation will capture all relevant ADMET signals equally well.

The broader implication is that ADMET benchmarking should not be treated only as a contest between universal molecular encoders. In the TDC ADMET group, the task identity itself carries important information: tasks differ in assay mechanism, dataset size, metric and relevant chemical signal. This view is consistent with systematic studies of molecular property prediction, where learned representations do not uniformly outperform fixed representations and where evaluation design and dataset structure can change the apparent winner [30, 7]. A practical ADMET system may therefore need to report the model setup used for each task, rather than a single architecture name for the entire benchmark.

The ablation tests clarify why this matters. Chemical descriptor models gave the largest contribution, showing that classical cheminformatics features remain useful even when deep molecular encoders are available. EPT/3D features, prediction averaging, seed averaging and task-wise model choice also contributed. The contribution is therefore not a new universal encoder, but a task-wise system for combining the chemical views that matter for each ADMET task.

The case studies serve a narrower but useful purpose. They do not validate mechanistic pharmacophores, but check whether task-specific predictors behave sensibly when the analysis moves from benchmark-level rankings to individual chemical decisions. Such a molecule-level view is important because practical ADMET decisions are made on individual compounds rather than benchmark averages. The AMES example links a high predicted mutagenicity score to a known high-risk motif class. The Deutivacaftor example links an external FDA-label compound to a transporter task and shows sensitivity to a local polar linker.

Reproducibility is especially important for a task-wise framework. Because the framework contains many candidate models, the final model setup for each task must be reported clearly. In this study, model type, averaging rule and seed set were recorded before official test labels were used. This separation is a benchmarking safeguard, not the main methodological contribution; it prevents repeated test-set checking from becoming hidden test-set selection [11, 7].

Several limitations remain. First, a task-wise system is harder to document and reproduce than a single neural backbone, because each task can use a different predictor, seed set or averaging rule. Second, averaged predictors are more expensive to store and run than a single model, which matters if the system is used for large virtual screens. Third, the framework cannot create assay signal that is absent from the data. Negative-margin tasks such as bioavailability, lipophilicity, VDss, DILI, hERG and LD50 are likely limited by a mixture of endpoint noise, chemical coverage, label uncertainty and strong public baselines; simply adding another selector is unlikely to solve them. Fourth, the molecule-level examples are behavioral checks, not mechanistic proof. Perturbation tests based on reconstructed or proxy models do not provide exact attribution for the whole final model and do not establish causal pharmacophores [14].

Future work should therefore focus less on adding another generic backbone and more on improving the evidence available for difficult endpoints. Useful directions include larger external test sets, prospective assays for negative-margin tasks, calibrated uncertainty estimates, endpoint-specific metadata such as assay conditions, and automated task agents that can choose candidate models, flag weak evidence and request additional experimental data before a prediction is treated as reliable [10].

## Conclusion

Trimole-Hybrid treats ADMET prediction as a task-specific modeling problem rather than a search for one universal molecular representation. On the 22-task TDC ADMET benchmark, the final model setups matched or exceeded the public TDC top-1 reference on 10 tasks and ranked in the top 10 on 21 tasks. Ablation tests showed that chemical descriptor models, EPT/3D features, prediction averaging, seed averaging and task-wise model choice each contributed to the final result. For benchmark users, the practical implication is clear: ADMET reports should record the molecular representation and model combination used for each task.

## Funding

This study was supported by the Shenzhen Medical Research Fund (award no. A2503002 to Y.L.), the National Key R&D Program of China (grant no.2025YFA0923500 to Y.L.), the Chinese University of Hong Kong (CUHK; award nos. 4937025, 4937026, 5501517, 5501329 and SHIAE BME-p1-24 to Y.L.), and the IdeaBooster Fund (award nos. IDBF23ENG05 and IDBF24ENG06 to Y.L.) and partially supported by grants from the Research Grants Council of the Hong Kong Special Administrative Region (Hong Kong SAR), China (project nos. CUHK 24204023 and 14208525 to Y.L.), and grants from the Innovation and Technology Commission of the Hong Kong SAR, China (project nos. GHP/065/21SZ, ITS/247/23FP and PRP/033/24FX to Y.L.). This research was also supported by the Research Matching Grant Scheme at CUHK (award nos. 8601603 and 8601663 to Y.L.)

## Notes

https://github.com/dchen0212/trimole_hybrid

### Competing Interest Statement

The authors have declared no competing interest.

