## Supplementary Information for "A multimodal representation learning platform for accurate molecular ADMET prediction"

---

This file contains Supplementary Notes 1–7, Supplementary Figure S1 and Supplementary Tables A–C, S1–S18 and S4e for the Trimole-Hybrid ADMET study.

Repository: https://github.com/dchen0212/trimole_hybrid

### Contents

**Supplementary Overview** **2**

**Supplementary Note 1: Benchmark Protocol and Frozen Reference Snapshot** **2**

**Supplementary Note 2: Candidate Pool and Task Configuration Records** **2**

**Supplementary Note 3: Model Formulation and Task Configuration Matrix** **3**

**Supplementary Note 4: Formal Ablation Protocol** **4**

**Supplementary Note 5: Molecule-Level Case-Study Audit** **6**

**Supplementary Note 6: Metric Direction and Endpoint Scale** **6**

**Supplementary Note 7: Software Environment and Data Availability** **7**

**Supplementary Tables** **7**

### Supplementary Overview

This Supplementary Information provides the benchmark protocol, reference snapshot, task-configuration records, ablation settings, case-study checks, metric interpretation and software environment for the Trimole-Hybrid ADMET study. It does not add leaderboard claims beyond the main text. Complete machine-readable tables are provided in the GitHub repository under supplementary_tables/, including Supplementary_Tables_S1_S18.xlsx and the individual CSV files.

Supplementary Tables A–C are compact tables included in this PDF for readability. Supplementary Tables S1–S18 and S4e are the machine-readable workbook and CSV tables used as the complete record for the benchmark, ablations, case-study checks and provenance information.

**Supplementary file guide.** Supplementary Tables A–C and Supplementary Figure S1 provide compact PDF summaries. Supplementary Tables S1–S18 and S4e provide the complete machine-readable record. Table S1 reports the full 22-task benchmark; Tables S2, S13 and S15 record task configurations and selection evidence; Tables S3 and S14 record the fixed public-reference snapshot; Tables S4 and S4e record ablation and naive MLP late-fusion control results; Tables S7–S10 and S17 record molecule-level case-study checks; Table S11 records split and source provenance; Table S16 records metric direction and endpoint-scale interpretation; and Table S18 records the software environment.

### Supplementary Note 1: Benchmark Protocol and Frozen Reference Snapshot

All benchmark results used the 22 tasks in the TDC ADMET benchmark group and the official scaffold splits. For each task, the official training and validation sets were used for model fitting, model choice and calibration. The official test set was evaluated only after the task-specific model configuration had been fixed.

The public TDC leaderboard is not a set of baselines rerun by us under one budget. We therefore used a fixed leaderboard snapshot recorded on 15 May 2026. The top-1 reference score, task rank and metric direction were copied from that snapshot and treated as external comparison metadata. These values were not used to choose model families, blend weights, seed sets, ablation variants or molecule-level examples.

A positive direction-normalized margin means that Trimole-Hybrid matched or exceeded the fixed public top-1 score after accounting for metric direction. This comparison should not be read as a head-to-head rerun of every public baseline.

Supplementary Table S14 records the snapshot date, the public reference method name when available, the reference score, the Trimole-Hybrid score, the direction-normalized margin and the rank metadata.

### Supplementary Note 2: Candidate Pool and Task Configuration Records

For each task, Trimole-Hybrid selected one model configuration from a fixed candidate pool. The pool included sequence-based encoders, graph-based encoders, EPT/3D predictors, chemistry-prior models, seed-bagged variants and prediction-level ensembles. Selection used only the official training/validation split or scaffold cross-validation within the training/validation partition.

The selected-configuration ledger records the model family, task configuration, seed usage, ensemble rule, EPT/3D use and source files. Supplementary Table A provides a compact version of that ledger. Full candidate identifiers, weights, seed definitions, source files and ablation configurations are given in Supplementary Tables S2–S2d. Supplementary Tables S13 and S15 give the number of checked variants and the candidate-pool scope for each task.

### Supplementary Note 3: Model Formulation and Task Configuration Matrix

The main Methods section is short to meet the Bioinformatics length constraint. Here we give the notation used for implementation. For task $t$, let $D_t^{\text{train}}$, $D_t^{\text{val}}$ and $D_t^{\text{test}}$ denote the official training, validation and test splits. Model development used

$$D_t^{\text{dev}} = D_t^{\text{train}} \cup D_t^{\text{val}}, \tag{1}$$

whereas $D_t^{\text{test}}$ was used only after the task-specific configuration had been fixed. The same rule was used for model-family choice, blend-weight choice, seed choice, case-study promotion and ablation design.

For molecule $x_i$, the candidate models used the following molecular representations:

$$\mathcal{V}(x_i) = \{v_{\text{seq}}(x_i), v_{\text{graph}}(x_i), v_{\text{3D}}(x_i), v_{\text{chem}}(x_i)\}, \tag{2}$$

where the four terms denote sequence, molecular graph, geometry-sensitive and chemistry-prior representations, respectively. Each available representation is mapped to an embedding or feature vector,

$$h_i^{(m)} = f_m(v_m(x_i)), \qquad m \in \mathcal{M}_t, \tag{3}$$

where $\mathcal{M}_t$ is the modality set available for task $t$. Sequence candidates used canonical SMILES language-model representations. Graph candidates used pretrained molecular graph embeddings. EPT/3D candidates used geometry-sensitive or task-tailored structural information. Chemistry-prior candidates used fingerprints, descriptors and tabular chemical features.

For neural fusion candidates, modality-specific embeddings were projected into a shared hidden space,

$$z_i^{(m)} = \phi_m\left(h_i^{(m)}\right). \tag{4}$$

In gated fusion variants, projected embeddings were combined as

$$h_i^{\text{fuse}} = \sum_{m \in \mathcal{M}_t} \alpha_{i,t}^{(m)} z_i^{(m)}, \qquad \sum_{m \in \mathcal{M}_t} \alpha_{i,t}^{(m)} = 1, \tag{5}$$

where $\alpha_{i,t}^{(m)}$ denotes a sample- or task-dependent modality weight. MLP-fusion variants concatenated projected embeddings instead of applying gates. The task-specific head then produced

$$\hat{y}_{i,t} = g_t\big(h_i^{\text{fuse}}\big), \tag{6}$$

with $g_t$ defined as a classification or regression head according to the official TDC task type.

Some candidates combined predictions from fixed base models rather than changing the molecular encoders. For task $t$, an ensemble candidate can be written as

$$\hat{y}_{i,t}^{\text{ens}} = \sum_{k=1}^{K_t} w_{k,t}\, T_t\left(\hat{y}_{i,t}^{(k)}\right), \qquad \sum_{k=1}^{K_t} w_{k,t} = 1, \tag{7}$$

where $\hat{y}_{i,t}^{(k)}$ is the prediction from candidate $k$, $w_{k,t}$ is a validation-selected or predefined ensemble weight, and $T_t(\cdot)$ is a task-dependent transformation such as identity, logit, rank or z-score transformation. Identity transformations were used for raw regression-value blending. Rank- or z-score-based transformations were used for Spearman-style tasks.

Task-specific selection used validation data only. Let $\mathcal{C}_t$ denote the candidate set for task $t$. The selected configuration was

$$c_t^* = \arg\max_{c \in \mathcal{C}_t} S_t^{\text{val}}\big(c; D_t^{\text{val}}\big), \tag{8}$$

where $S_t^{\text{val}}$ is the official task metric after applying the appropriate metric direction. After selection, $c_t^*$ was fixed and evaluated on the official test split to obtain the final reported score,

$$S_t^{\text{test}} = S_t\big(c_t^*; D_t^{\text{test}}\big). \tag{9}$$

The same validation-only rule was used for the final model and for the component-removal ablations.

**Supplementary Table A.** Compact component matrix for the fixed 22-task model configurations. A filled entry indicates that the component was used in the final configuration for that task. Full configuration records are provided in Supplementary Tables S2–S2d.

| Dataset | Metric | Fixed model family | Chem | Seq | Graph | EPT/3D | Blend | Bag |
|---|---|---|---|---|---|---|---|---|
| TDC.Bioav | AUROC | Fixed chem ExtraTrees | ● | | ● | | | ● |
| TDC.Caco2 | MAE | Fixed chem ExtraTrees | ● | ● | ● | | | ● |
| TDC.HIA | AUROC | Task-specific chem blend | ● | | | | ● | ● |
| TDC.Lipo | MAE | XL/chem hybrid blend | ● | ● | ● | ● | ● | ● |
| TDC.Pgp | AUROC | KPGT/EPT logit blend | | ● | ● | ● | ● | |
| TDC.AqSol | MAE | FP-XGB/KPGT-EPT/XL blend | ● | ● | ● | ● | ● | |
| TDC.BBB | AUROC | Descriptor ExtraTrees | ● | | | | | |
| TDC.PPBR | MAE | KPGT/fingerprint blend | ● | | ● | | ● | ● |
| TDC.VD | Spearman | KPGT/EPT rank blend | ● | | ● | ● | ● | ● |
| TDC.CYP2C9-S | AUPRC | Fixed chem XGBoost | ● | ● | ● | ● | | ● |
| TDC.CYP2C9-I | AUPRC | Fixed chem XGBoost | ● | | ● | ● | | ● |
| TDC.CYP2D6-S | AUPRC | AP/XL/descriptor blend | ● | ● | | ● | ● | ● |
| TDC.CYP2D6-I | AUPRC | KPGT/EPT/FP rank blend | ● | | ● | ● | ● | ● |
| TDC.CYP3A4-S | AUROC | Family-transfer FP-XGBoost | ● | | | | | |
| TDC.CYP3A4-I | AUPRC | XL/KPGT-EPT logit blend | ● | ● | ● | ● | ● | ● |
| TDC.CL-Hepa | Spearman | Family-transfer FP-XGBoost | ● | | | | | |
| TDC.CL-Micro | Spearman | KPGT/EPT seedbag | | | ● | ● | ● | ● |
| TDC.Half_Life | Spearman | C+K descriptor/z-score blend | ● | ● | ● | | ● | ● |
| TDC.AMES | AUROC | Repeated fingerprint XGBoost | ● | | | | | ● |
| TDC.DILI | AUROC | XL/KPGT-EPT raw blend | ● | ● | ● | ● | ● | ● |
| TDC.hERG | AUROC | Fixed chem ExtraTrees | ● | ● | | ● | | ● |
| TDC.LD50 | MAE | Fixed chem XGBoost | ● | ● | ● | ● | | ● |

The source-provenance check found that the v36 main results used official TDC benchmark splits and did not use `data_new`. Supplementary Table S11 gives the main-result source check. The ablation README records the same rule: fixed candidate families were rerun on official-split embeddings or fingerprints, validation data were used for selection, and test scores were used only after selection.

Supplementary Table S13 is a one-row-per-task manifest. It links the final score, selected candidate, model-family summary, seed definition, checked-variant count, candidate-pool scope, selection summary, test-score source file, official-split provenance and component flags. Supplementary Table S15 summarizes the validation-only selection record for each task and points to the selected configuration and the five component-removal comparators.

### Supplementary Note 4: Formal Ablation Protocol

The ablations quantified the contribution of each major component to the final benchmark scores. The variants were: full Trimole-Hybrid, no task-adaptive selection, no chemistry sidecar, no EPT/3D, no prediction-level ensemble and a single-seed version of seed-bagged tasks where available.

For each component-removal ablation, task-wise selection was repeated with the same validation-only rule after removing that component class. The single-seed variant reports the closest available canonical single-seed candidate. These variants were not submitted as independent leaderboard entries, so they should be read as within-study comparisons against the full model. The ablation source files also record that the reruns used official-split embeddings, fingerprints and fixed heads, and that `data_new` was not used.

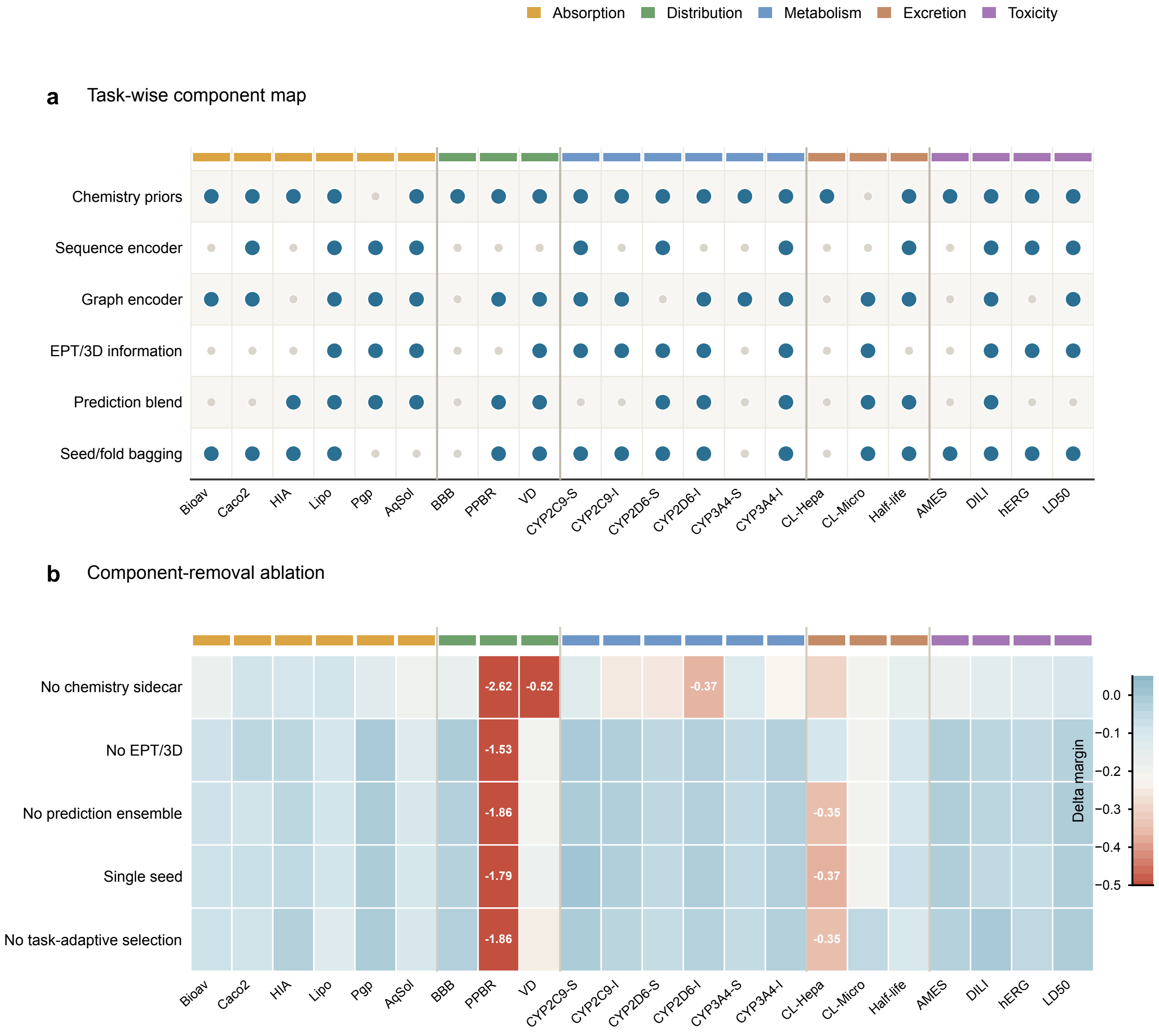


**Supplementary Figure S1:** Task-wise component usage and component-removal ablation effects. **a**, Component map for the 22 fixed TDC ADMET task configurations. Blue circles indicate components used in the final configuration; grey circles indicate components not used. **b**, Direction-normalized margin changes after removing each component class relative to the full v36 configuration. Red indicates loss relative to the full configuration. Values are clipped at −0.5 for readability; exact values are provided in Supplementary Table S4. Task labels use dataset codes; full endpoint names are listed in Supplementary Table S1.

### Supplementary Note 5: Molecule-Level Case-Study Audit

The main text contains two molecule-level sensitivity examples. Both were chosen after the task models had been fixed. They were not used for model selection, task promotion, ablation design or leaderboard comparison. They report score changes for individual molecules and matched local replacements. They do not prove causal pharmacophores.

The two examples have different replay status. The AMES example uses the saved full prediction for the original official-test molecule, but the perturbation sweep used a reconstructed AMES task model. We therefore label it as reconstruction-based sensitivity, not exact full-ensemble attribution. The Deutivacaftor/P-gp example uses the replayed P-gp pipeline for both the external molecule and its matched local replacement. The P-gp perturbation therefore has the stronger replay status. Both examples remain model-sensitivity checks, not mechanistic claims.

**Supplementary Table B.** Two molecule-level sensitivity examples shown in Figure 4. Full details are provided in Supplementary Table S7.

| Case | Task | Label | MiniMol score | Trimole score | Sensitivity status |
|---|---|---|---|---|---|
| AMES test_297 | AMES | 1 | 0.293194 | 0.894085 | Original molecule uses a saved Trimole-Hybrid prediction; perturbation uses a reconstructed AMES model and is not exact full-ensemble attribution. |
| Deutivacaftor | P-gp | 1 | 0.746979 ± 0.064799 | 0.914545 ± 0.021395 | Exact replayed P-gp pipeline; O-to-CH2 matched replacement reduced the score to 0.729853 ± 0.149185. |

External molecule candidates were converted to canonical SMILES and compared with the corresponding official TDC task splits. Candidates found in the training set were excluded from main-text use. For Deutivacaftor/P-gp, no canonical SMILES overlap was found in the P-gp training, validation or test split. This example is therefore treated as an external molecule-level check, not as a new benchmark task.

Several molecule-level candidates were examined but not used in the main sensitivity figure. A candidate was not promoted when an exact replay path was not available under the archived inference setting, the matched replacement did not yield a stable sensitivity pattern, or the Trimole-Hybrid prediction did not clearly improve on the MiniMol-style baseline. Supplementary Table S17 lists the promoted and non-promoted molecule-level examples. A case was eligible for main-text use only if it had a clear label or official-test label, no prohibited training-set overlap for external cases, a MiniMol-style comparison where relevant, a reportable Trimole-Hybrid prediction and a replay-status label.

For this reason, AMES is listed in the Figure 4 table but labeled as reconstruction-based sensitivity, whereas Deutivacaftor/P-gp is labeled as exact replay. The two evidence levels should not be treated as equivalent.

### Supplementary Note 6: Metric Direction and Endpoint Scale

The benchmark uses classification metrics, absolute-error regression metrics and rank-correlation metrics. Direction-normalized margins were used only to compare these task-level scores in one table. For AUROC, AUPRC and Spearman correlation, the margin is the Trimole-Hybrid score minus the fixed reference score. For MAE, the margin is the reference MAE minus the Trimole-Hybrid MAE. Positive margins therefore indicate matched or better performance relative to the fixed reference.

Supplementary Table S16 records the metric direction, margin formula and label-scale notes used in the manuscript. AUROC, AUPRC and Spearman correlation are task-level metrics. A single molecule can have a predicted classification score, and a single MAE-task molecule can have an absolute error on the reported label

**Supplementary Table C.** Selected case-study candidates that were not used in the main figure. Full details are provided in Supplementary Table S10.

| Candidate | Task | Main issue | Final use |
|---|---|---|---|
| BBB external candidates | BBB | The archived replay path did not meet the criteria for main-text interpretation, and no MiniMol-wrong/Trimole-correct external candidate was found. | Not used in the main figure. |
| Ritlecitinib | CYP3A4-S | Original prediction comparison was promising, but matched replacement did not give a stable sensitivity signal. | Not used in the main figure. |
| Mavorixafor | P-gp | Boundary case with weak replacement evidence. | Not promoted. |
| AqSol external candidates | AqSol | Exact arbitrary-SMILES external inference was not available in the archived replay setting. | Not promoted. |
| PPBR external candidates | PPBR | External predictions remained near-reconstruction or proxy rather than exact replay. | Not promoted. |

scale. A single molecule cannot have its own AUROC, AUPRC or Spearman value. Spearman-task molecule examples should therefore be read as continuous-score examples, not as single-molecule rank-correlation errors.

### Supplementary Note 7: Software Environment and Data Availability

Supplementary Table S18 reports the software environments used for model assembly, official-split tabular models and graph-feature extraction. The public project repository is available at https://github.com/dchen0212/trimole_hybrid. The principal benchmark environment used Python 3.10.20 with RDKit 2026.03.1, scikit-learn 1.7.2, XGBoost 3.2.0 and PyTorch 2.5.1 with CUDA 12.1. KPGT feature extraction used a separate Python 3.10.19 environment with DGL 2.0.0 and PyTorch 2.1.2 with CUDA 12.1.

We include this environment table because some exploratory replay analyses depended on archived model files and package versions. AMES remains a valid benchmark result, but its perturbation result is labeled as reconstruction-based because the exact archived full-ensemble replay path was not available for arbitrary AMES SMILES.

All supplementary tables are included in the manuscript package under `supplementary_tables/`. Source code, supplementary tables and lightweight audit manifests are available from the main manuscript repository. Large trained model binaries, cached embeddings and serialized estimators will be provided through archive or release assets where storage and licensing permit. The supplementary workbook remains the manuscript-level record for the benchmark, ablations and case-study checks.

### Supplementary Tables

**Supplementary Table Inventory.** Complete machine-readable tables are provided in the supplementary workbook and as individual CSV files.

| Table | File | Description |
|---|---|---|
| S1 | `Table_S1_TDC_ADMET22_benchmark.csv` | Full 22-task benchmark table, including category, dataset, metric, direction, Trimole-Hybrid score, fixed public top-1 score, margin, rank and task configuration. |

| Table | File | Description |
|---|---|---|
| S2 | Table_S2_endpoint_recipe_and_variant_ledger.csv | Task-wise model configurations and ablation variants, including selected candidates, source files and component-use flags. Additional files Table_S2c_final_endpoint_summary.csv and Table_S2d_endpoint_overview_for_pdf.csv provide compact final-task summaries. |
| S3 | Table_S3_frozen_reference_snapshot.csv | Fixed reference snapshot and matched multibaseline data used for benchmark interpretation and Figure 2 support. |
| S4 | Table_S4_formal_ablation_long.csv | Formal ablation scores, margins, deltas and heatmap-ready data for all 22 tasks and ablation variants. The naive MLP late-fusion control is provided separately in Table_S4e_naive_mlp_late_fusion_control.csv. |
| S4e | Table_S4e_naive_mlp_late_fusion_control.csv | Naive MLP late-fusion control scores and losses relative to the final fixed task setups, supporting Figure 3e. |
| S5 | Table_S5_ADMET_category_summary.csv | ADMET-category and metric-family summaries used to support Figure 2. |
| S6 | Table_S6_extended_official_test_case_examples.csv | Extended official-test molecule-level examples retained as supplementary material. |
| S7 | Table_S7_Figure4_two_case_sensitivity_audit.csv | Figure 4 two-case table, including labels, prediction scores, perturbations, replay status and interpretation scope. |
| S8 | Table_S8_external_candidate_leakage_audit.csv | External molecule leakage audit and P-gp split-overlap summary. |
| S9 | Table_S9_Pgp_matched_replacement_sweep.csv | P-gp matched-replacement sweep and exact replay audit supporting the Deutivacaftor/P-gp sensitivity example. |
| S10 | Table_S10_not_promoted_case_audit.csv | Explored molecule-level candidates that were not used in the main figure, with the reason for exclusion. |
| S11 | Table_S11_split_and_source_provenance_audit.csv | Official-split and source-provenance check for the main v36 benchmark results. |
| S12 | Table_S12_external_case_alignment_feasibility.csv | External case alignment feasibility table and selected external search snapshot. |
| S13 | Table_S13_reproducibility_manifest.csv | One-row-per-task manifest linking selected model, seed definition, selection summary, official-split provenance, test-score source file and component flags. |
| S14 | Table_S14_frozen_reference_snapshot_audit.csv | Fixed TDC reference snapshot, including the snapshot date, public reference metadata where available, Trimole-Hybrid score, reference score, margin and rank metadata. |
| S15 | Table_S15_validation_only_selection_evidence.csv | Validation-only selection summary for all 22 tasks, including checked-variant count, candidate-pool scope and candidate notes. |
| S16 | Table_S16_metric_direction_and_unit_guide.csv | Metric direction, margin formula, endpoint-scale and single-molecule interpretation guide. |
| S17 | Table_S17_case_study_decision_boundary.csv | Case-study decision table for promoted and non-promoted molecule-level examples. |
| S18 | Table_S18_software_environment_manifest.csv | Server-side software and execution-environment manifest. |
